# Capacity of short-term memory in dyslexia is reduced due to less efficient utilization of items’ long-term frequency

**DOI:** 10.1101/2020.03.25.008169

**Authors:** Eva Kimel, Itay Lieder, Merav Ahissar

## Abstract

Dyslexia, defined as a specific impairment in decoding the written script, is the most widespread learning difficulty. However, individuals with dyslexia (IDDs) also consistently manifest reduced short-term memory (STM) capacity, typically measured by Digit Span or non-word repetition tasks. In this paper we report two experiments which test the effect of item frequency and the effect of a repeated sequence on the performance in STM tasks in good readers and in IDDs. IDDs’ performance benefited less from item frequency, revealing poor use of long-term single item statistics. This pattern suggests that the amply reported shorter verbal spans in dyslexia may in fact reflect their impaired sensitivity to items’ long-term frequency. For repeated sequence learning, we found no significant deficit among IDDs, even when a sensitive paradigm and a robust measure were used.

---

Developmental dyslexia is the most widespread learning difficulty, defined as a specific difficulty in accurate or fluent word recognition, decoding, and spelling abilities (American Psychiatric Association, 2013) in spite of adequate hearing levels, normal intelligence and adequate educational opportunities. Individuals with dyslexia (IDDs) are characterized, beyond their reading difficulties, as having poor short term memory (STM) (e.g. Jeffries & Everatt).

STM of IDDs is typically assessed with the standard Digit Span test (e.g. Snowling, 2000; Spring, 1976). Performance in Digit Span is correlated with reading speed, within and across languages (Naveh-Benjamin & Ayres, 1986), suggesting functional relations between poor span scores, reflecting poor STM, and poor reading skills among IDDs. Using repetition of non-words or lists of non-words also yields poor STM scores among IDDs (e.g. Jeffries & Everatt, 2004; S Roodenrys & Stokes, 2001; M. Snowling, Goulandris, Bowlby, & Howell, 1986; M. J. Snowling, 1981). Overall, the observation of poor scores in span tasks among IDDs is very robust (though see Wimmer, 1993). The impaired performance in STM tasks among IDDs was traditionally attributed to poor phonological representations (e.g. Snowling, 2000), or to reduced efficiency in accessing these representations (Ramus, 2014; Ramus & Szenkovits, 2008).

However, the hypothesis that the core deficit in dyslexia stems from poor phonological representations was challenged by many studies in the past two decades, showing that IDDs have non-phonological deficits, and have adequate performance in some tasks that require adequate phonological representations (Ramus & Ahissar, 2012). One such hypothesis (the anchoring deficit hypothesis, Ahissar, 2007; Ahissar, Lubin, Putter-Katz, & Banai, 2006; Oganian & Ahissar, 2012), relates to IDDs’ learning patterns and proposes that their process of automatic learning of statistics in the input is impaired (for a Bayesian account of the anchoring deficit, see Jaffe-Dax, Raviv, Jacoby, Loewenstein, & Ahissar, 2015). Specifically, it proposes that IDDs’ implicit memory trace decays faster than that of good readers (Jaffe-Dax, Kimel, & Ahissar, 2018; Lieder et al., 2019), consequently reducing their learning slope as a function of exposure, and impeding the development of refined representations of highly frequent stimuli, including, but not specific to phonological stimuli (Banai & Ahissar, 2017; Jaffe-Dax, Lieder, Biron, & Ahissar, 2016). One prediction of this hypothesis is that IDDs will also underuse morphological representations, which heavily rely on linguistic regularities. Indeed, it was recently shown that IDDs rely less on morphology when acquiring new vocabulary, suggesting a novel account to IDDs’ reduced vocabulary (Kimel & Ahissar, 2019).

Span tasks provide an informative window into the use of long-term statistics, since repeated (long-term) exposure has a large effect on the performance of the general population in these tasks. Thus, spans of frequent words are larger than those of infrequent ones (Hulme et al., 1997; Roodenrys, Hulme, Lethbridge, Hinton, & Nimmo, 2002), the span of words is larger than the span of non-words (Hulme, Maughan, & Brown, 1991), repetition accuracy of a multi-syllabic non-word is higher when the frequency of the first syllable is high (Tremblay, Deschamps, Baroni, & Hasson, 2016), and recall of sequences of syllables are higher for syllables that occur frequently in polysyllabic English words (Nimmo & Roodenrys, 2002).

The (long-term) frequency effect in span tasks provides an opportunity to test theories of the mechanisms underlying dyslexia. Theories that view IDDs’ phonological deficits as a cause rather than an accumulative outcome of learning deficits, predict that novel challenging phonological representations will pose greater difficulties to IDDs than familiar stimuli, since massive exposure provides an opportunity to compensate for the impaired phonology.

However, the anchoring hypothesis proposes that IDDs will not benefit from item frequency to the same extent as controls, and therefore their relative difficulties will increase with item frequency. Namely, exposure will facilitate performance of all individuals, but it will have a smaller effect on that of IDDs. Hence, group effect will increase with item frequency, in agreement with theories that focus on slower learning rate (e.g. Gabay, Thiessen, & Holt, 2015; Nicolson, Fawcett, Brookes, & Needle, 2010; Roderick I. Nicolson & Fawcett, 2007), rather than on reduced phonological skills as the core mechanism underlying dyslexia.

## Experiment 1: Digit Span

We designed Experiment 1 to test this hypothesis. We used the most common assessment of IDDs’ poor STM – the standard Digit Span task (e.g. (Ackerman, Dykman, & Gardner, 1990; Torgesen, Wagner, Simmons, & Laughon, 1990). In order to compare the span of the frequent digits to infrequent items with the same semantic priors (which could assist word recall), we compared Digit Spans in participants’ native language, Hebrew, with their Digit Span in a foreign (though familiar) language, English. All Hebrew speaking participants were highly familiar with English, yet they had substantially reduced exposure to digits spoken in English throughout their lives. We hypothesized that IDDs would gain less from the frequent exposure to Hebrew digits compared with English digits, and therefore group difference will be smaller for English digits.

This prediction is counter-intuitive, since IDDs are known to have difficulties in second language (Crombie, 1997; Ganschow, Sparks, & Schmeider, 1995; Sparks & Ganschow, 1991; Spolsky, 1989). Moreover, according to the phonological deficit hypothesis (e.g. M. J. Snowling, 2000) foreign language words are expected to be harder to process by IDDs, since foreign language phonology is expected to pose a bigger phonological challenge. We do not dispute the difficulty of IDDs in acquiring a second language, but we explain their difficulty as a consequence of a slower learning curve of the trained items. Therefore, the relative difficulty of IDDs should increase with exposure, since their item-specific long-term learning slope is shallower. Thus, taking learning into account yields a unique prediction.

## Method

### Cognitive assessments

1. The Block Design task (a subtest from the Hebrew version of the Wechsler Adult Intelligence Scale, WAIS-III; Wechsler, 1997) was used as a measure of non-verbal intelligence. This task is often used to match groups for non-verbal reasoning.
2. The Digit Span task (Wechsler, 1997), a standard STM measure.

### Reading and phonological measures

1. Single word reading: words, pseudo-words and non-words. The lists of real words and pseudo-words were standard lists (Deutsch & Bentin, 1996). The non-words did not follow Hebrew morphology or phonotactics and were taken from the Aztec language Nahuatl (Oganian & Ahissar, 2012).
2. Paragraph reading (Ben-Yehudah, Sackett, Malchi-Ginzberg, & Ahissar, 2001).
3. Phonological awareness: 20 word pairs were orally presented (Ben-Yehudah & Ahissar, 2004; Ben-Yehudah et al., 2001). For each pair, participants were asked to swap the initial phonemes of the two words (the “spoonerism” task).

### Participants

We recruited 37 adult participants with dyslexia and 35 age-matched controls through ads posted at the Hebrew University campuses, and in two other colleges in Jerusalem. Participants were paid for their participation, and took part also in other studies at the lab (e.g. Kimel & Ahissar, 2019). All participants received all their schooling in Israel.

Exclusion criteria included psychiatric medications (other than attention deficit medications), hearing problems, extensive musical background and below average cognitive scores (for details see Kimel & Ahissar, 2019). Based on these criteria, 1 control participant and 1 dyslexic participant were excluded from the study. One additional IDD had accurate (100% correct) non-word reading, and was therefore excluded from the study. Additionally, data of one control participant were excluded in order to improve age matching between the groups (prior to data analysis). This process resulted in 35 IDDs and 33 control participants matched for age and general reasoning skills, reported in Table 1. Due to technical reason, 32 IDDs and 30 controls participated in Experiment 1. Participants (IDDs, n=8), who routinely take medicine for ADHD (e.g., Concerta), did not take medicine on the days of the experiments. The performance of this sub-group did not differ from that of the other participants in any of the main effects in this study (the smallest p value for these comparisons was .41).

**Table 1.**
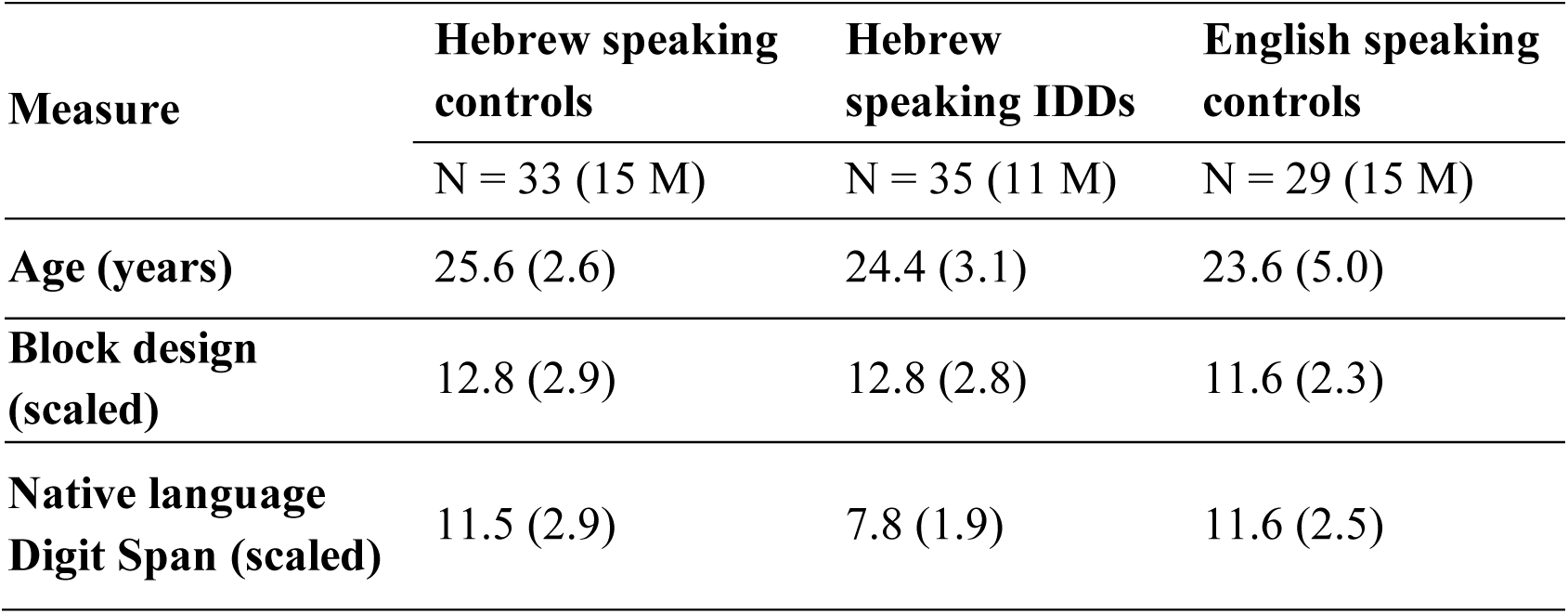
Age and standard cognitive scores (mean and SD) of the three groups: native Hebrew speaking controls, native Hebrew speaking IDDs, and native English speakers. The groups did not significantly differ in age and in block design. In line with previous reports, IDDs’ Digit Span scores were significantly lower than both control groups’ (Scheffe Post Hoc tests: Hebrew speaking controls vs. IDDs: d = 3.66, SE = .61, p < 10_-6_, Hebrew speaking IDDs vs. English speaking controls: d = 3.79, SE = .631, p < 10_-6_, Hebrew vs. English speaking controls: d = .14, SE = .64, p < .978). M – Male participants.

An additional group of participants was recruited to control for native Digit Span scores in English. We recruited group of native English speakers without reading difficulties through ads put at the school of international students of the Hebrew University. All recruited participants took Hebrew as a second language class, yet their Hebrew was rather basic. We applied similar exclusion criteria as for the Hebrew speakers (Kimel & Ahissar, 2019). Matching for age and cognitive scores (Block Design), resulted in a group of 29 English-speaking participants. Table 1 shows their general scores.

### Procedure

The Hebrew Digit Span was conducted according to the instructions in the Wechsler Adult Intelligence Scale for Digit Span forward (Wechsler, 1997). Participants were auditorily presented with sequences of digits and were requested to repeat the digits in the presented order. The experiment started with 2 sequences of 2 digits; sequence length increased by 1 digit after 2 sequences of the same length; the task continued until the participant either failed on 2 sequences of the same length or reached the maximal sequence length of 8 digits (Wechsler, 1997). Performance score was the number of correctly reproduced sequences (a measure frequently used in span tasks; e.g. Oganian & Ahissar, 2012).

The English Digit Span test was conducted using a digital recording of a native English speaker, the nine digits were recorded separately and then combined into sequences with increasing length. The sequences were played to the participants via headphones and their responses were recorded. Participants were requested to repeat each sequence immediately after it ended. An experimenter sat by the participant administered the task according to the Digit Span WAIS protocol. Assessments of Digit Span in the two languages were separated by other tasks. We asked the Hebrew speaking participants whether they translated digits presented in English to Hebrew and they reported that they did not. Hebrew proficiency of the English speaking participants was not high enough to perform Digit Span in Hebrew.

## Results

As expected, the group of IDDs had substantial difficulties in all reading and reading-related measures (Table 2). Consistent with previous findings, IDDs’ scores were significantly lower than those of controls in the Digit Span task.

**Table 2.** Means and SDs of the Hebrew speaking subjects with and without dyslexia in reading related tasks. M – Male participants.

| Measure | Controls | IDDs | Group difference (t) |
| --- | --- | --- | --- |
|  | N = 33 (15 M) | N = 35 (11 M) |  |
| Reading accuracy (% correct) |  |  |  |
| Single words | 97.1 (4.0) | 87.4 (7.8) | 6.3*** |
| Single pseudo-words | 90.8 (10.8) | 61.2 (18.1) | 8.0*** |
| Single non-words | 88.1 (12.9) | 51.1 (24.4) | 7.7*** |
| Paragraph reading | 98.7 (1.4) | 95.0 (4.1) | 4.7*** |
| Reading rate (words/minute) |  |  |  |
| Single words | 100.6 (32.7) | 69.9 (24.9) | 4.3** |
| Single pseudo-words | 60.4 (25.2) | 33.4 (11.7) | 5.6*** |
| Single non-words | 43.1 (15.5) | 25.7 (8.7) | 5.7*** |
| Paragraph reading | 140.5 (21.4) | 98.2 (22.5) | 7.8*** |
| Phonological Awareness: Spoonerism |  |  |  |
| Accuracy (% correct) | 92.1 (6.9) | 77.7 (17.0) | 4.5** |
| Rate (items/minute) | 10.1 (3.1) | 5.6 (3.0) | 6.1*** |

\*\* $p < .0001$
\*\*\* $p \leq .00001$

We first asked whether Digit Span carried out in a native language indeed produces an advantage. To test that, we compared the scores of Digit Span administered in English between English and Hebrew speakers. Indeed, Hebrew speaking controls had significantly lower scores in English (*t* = 2.89, *p <* .005; Figure 1A), even though the use of English digits is common among Hebrew speakers. Importantly, this difference was significant in spite of their matched scaled Digit Span scores in their respective native languages (Table 1). This difference established the case that span score indeed continue to increase even after massive exposure.

**Figure 1.**
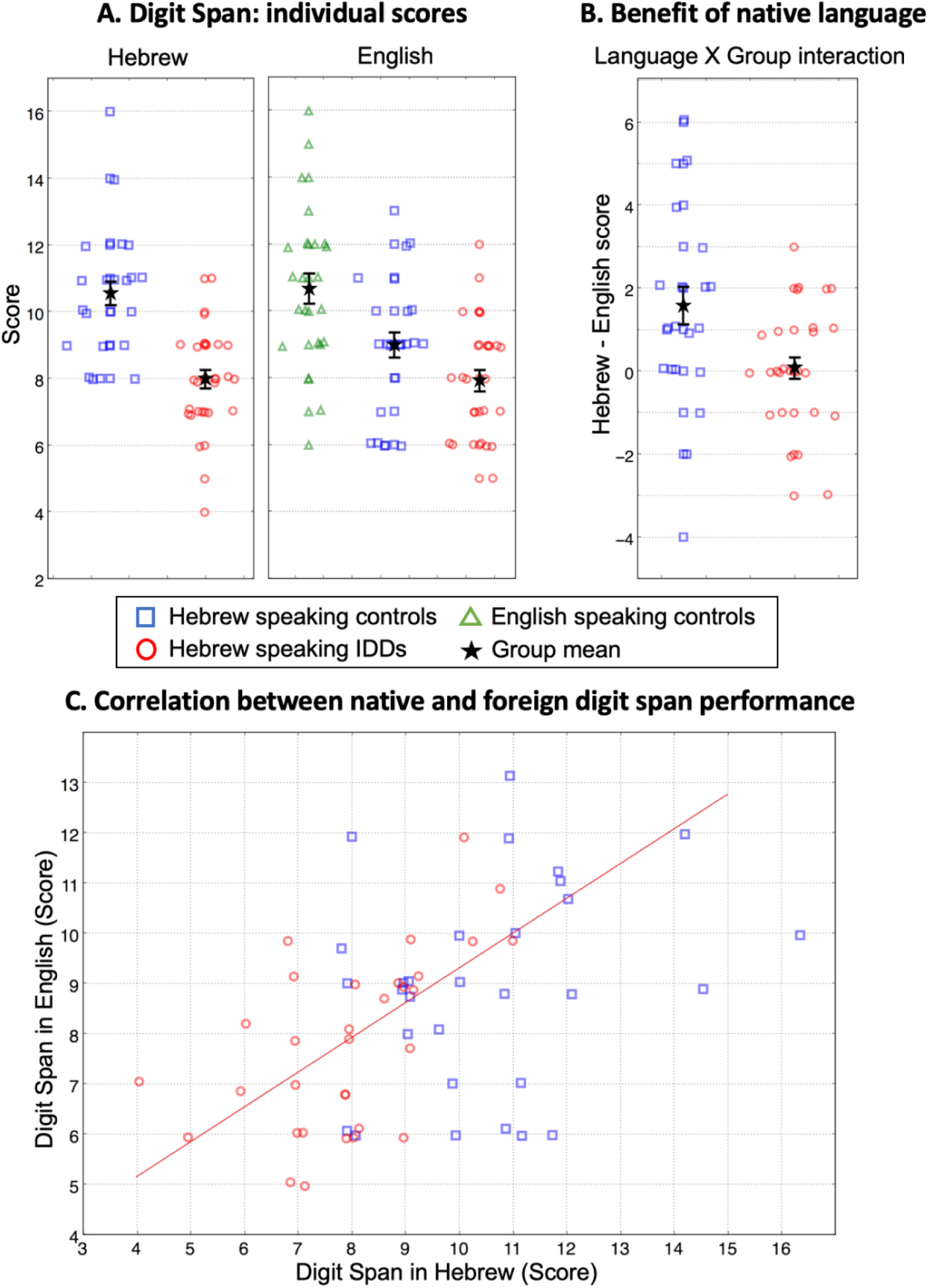
A. Left: Digit Span Scores of Hebrew speaking controls (blue squares) and IDDs (red circles) tested in Hebrew. Right: Scores of English speaking controls (green triangles), Hebrew speaking controls (blue squares), and IDDs (red circles) tested in English. B. Difference in scores between English and Hebrew, measuring the familiarity (long-term frequency) effect for digits. Hebrew speaking controls benefit more than Hebrew speaking IDDs from performing the span in Hebrew, their native language. Though Hebrew speaking IDDs’ raw scores are the same in English and in Hebrew, it reflects some benefit to Hebrew, in which digit names are typically di-syllabic and typical spans are shorter than in English (Naveh-Benjamin & Ayres, 1986). C. A scatter plot of Digit Span scores in English vs. scores in Hebrew. Scores of Hebrew speaking IDDs are highly correlated (Pearson: r = .60, p < .001) whereas those of Hebrew speaking controls are not (Pearson: r = .25, p = .179). Symbols denote individual scores. Error bars denote 1 SEM. The values are slightly jittered for display purposes.

To test the prediction that IDDs’ scores will be particularly impaired in high-frequency, native language items, we analyzed the number of correctly reproduced sequences in each condition (score) using a mixed-design analysis of variance (ANOVA), with *Language* (native, Hebrew vs. foreign, English) as a within-subject factor, and *Group* (Hebrew speaking controls vs. Hebrew speaking IDDs) as a between-subject factor. Though restricted by the interactions, overall scores of controls were higher than those of IDDs (main effect of *Group*: *F*_(1,60)_ = 21.38, *p <* 10_-4_, *η2 =*.263; Figure 1A), and higher for participants’ native language, Hebrew, compared with non-native English (main effect of *Language*: *F*_(1,60)_ = 10.06, *p <* .002, *η2 =*.144; Figure 1A).

As predicted, controls benefited significantly more than IDDs from performing Digit Span in their native language (*Group X Language*: *F*_(1,60)_ = 8.57, *p <* .005, *η2 =*.125; controls: *d =*1.57, *SE =*.37, *p <* 10_-4_, IDDs: *d =*.06, *SE =*.36, *p =* .862), and group difference was smaller in the infrequent (English) condition compared to the frequent (Hebrew) condition (native (Hebrew) language: *F*_(1,60)_ = 32.77, *p <* 10_-6_, *η2 =*.353, foreign (English) language: *F*_(1,60)_ = 4.71, *p <* .034, *η2 =*.073; Table 3; Figure 1B).

**Table 3.**
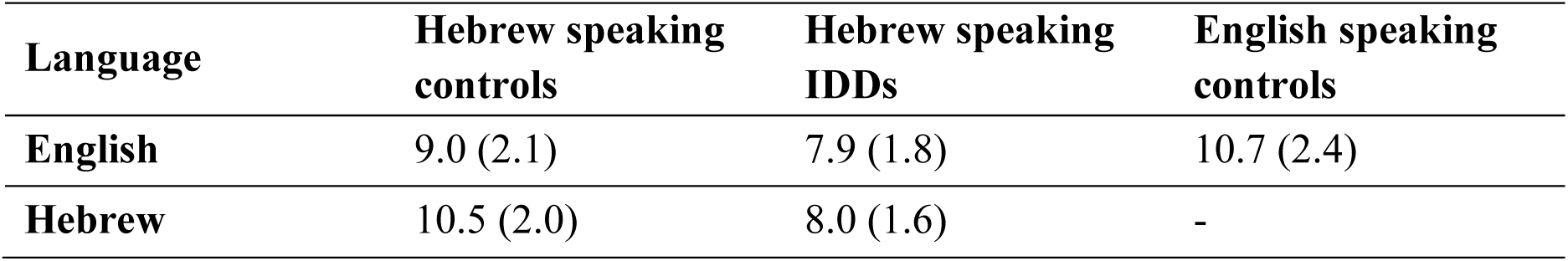
Experiment 1: scores (mean (SD)) of Digit Span forward, in Hebrew and English. The score for each sequence was either 0 or 1: 0 if there was at least one mistake/missing digit, and 1 if the recall was perfect. The total score for each participant for each condition is the sum of the sequences’ scores.

| **Language** | **Hebrew speaking controls** | **Hebrew speaking**<br>**IDDs** | **English speaking controls** |
| --- | --- | --- | --- |
| **English** | 9.0 (2.1) | 7.9 (1.8) | 10.7 (2.4) |
| **Hebrew** | 10.5 (2.0) | 8.0 (1.6) | - |

Given IDDs’ smaller gains when performing the Digit Span task in their native language, we reasoned that IDDs use the same set of mechanisms for both native and second language, relying less on statistical learning (Jaffe-Dax et al., 2015; Leider et al., 2019). We therefore hypothesized that their scores in native and in second language will be highly correlated. By contrast, controls gain substantially from long-term exposure. We hypothesized that for them, utilizing additional mechanisms of statistical learning underlie their elevated scores in native language, and thus correlation of their scores in first and second language will be reduced. Figure 1C illustrates that this was indeed the case: for IDDs the correlation between Digit Span scores in the two languages was highly significant (Pearson: *r =* .60, *p <* .001, Spearman *r =* .61, *p <* .001), whereas for controls it was not (Pearson: *r =* .25, *p =* .179, Spearman *r =* .27, *p =* .151; though the group difference between the strength of the correlations was only marginally significant: Fishers Z-test: one sided p < 0.05; Figure 1C).

## Summary and Conclusions

We administered Digit Span in Hebrew and in English to controls and IDDs whose native language is Hebrew. Controls benefited significantly more than IDDs from familiarity with digits presented in their native language. Hebrew speaking controls’ Digit Span in English was smaller than that of native English speakers, though their native-language Digit Span scores were matched, indicating that Hebrew speaking controls’ exposure to English is not sufficient for reaching asymptotic performance (even though English is heavily used in Israel).

As discussed above, the significant difference in performance of Digit Span between controls and IDDs was replicated many times. This observation was previously interpreted as reflecting generally limited verbal/phonological STM in dyslexia (e.g. Snowling, 2000; Spring, 1976). The results of Experiment 1 suggest a novel interpretation to IDDs’ poor spans. Namely, IDDs’ poor Digit Span reflects reduced accumulative, long-term implicit item-specific learning.

## Experiment 2: Hebb learning of serial order

Experiment 2 had two aims: 1. To ask whether the reduced effect of long-term frequency applies also to sub-lexical items – syllables. 2. To replicate previous studies, with mixed results, asking whether serial order learning is impaired in dyslexia.

Experiment 1 suggested that serial order repetition in dyslexia is substantially affected by long-term experience with the test items: group difference is bigger for items with which the participants have more experience. However, memory for order is also influenced by familiarity with sequences (i.e., series), and not just single items. Serial order learning was amply studied (e.g. Leclercq & Majerus, 2010; Reber, 1969, 1989; Saffran, Aslin, & Newport, 1996), and memory of repeated sequences has a great effect on STM. For example, it has been shown that frequent digit sequences specifically increase STM for digits (Jones & Macken, 2015, 2018), an effect, which cannot be solely explained by the frequency of the single digits.

Kimel et al. (Kimel, Weiss, Jakoby, Daikhin, & Ahissar, 2019), addressed benefit from item frequency and from repeated sequences in a paper studying syllable spans in four different populations: IDDs, controls, musicians and non-native speakers. They used the structure of the Digit Span task, but with frequent and infrequent syllables, and found that, as expected, individuals from the general population had substantially larger spans for frequent syllables compared with infrequent syllables. Musicians showed an enhanced effect of syllable frequency, and IDDs showed a reduced effect. Though consistent with the results of Experiment 1, these results for IDDs seem to contradict common assumptions and some previous findings.

First, though never directly tested for spans, previous reading studies suggested that IDDs have adequate benefit from long-term word frequency. Thus, they show the expected advantage from word frequency in reading (Davies, Cuetos, & Glez-Seijas, 2007; Van der Leij & Van Daal, 1999). We interpret this result as a consequence of a confound, when high-frequency words are used. Words both carry a meaning and have a phonological form, and thus, frequent words give a semantic and a phonological advantage. We derived a way to dissociate semantic from phonological frequency. In Experiment 1 we tested Digit Span in two languages, and thus semantic frequency was equated by design, in experiment 2 we use syllables, which typically lack a semantic component.

Kimel et al. (Kimel et al., 2019) also report that the four populations did not differ in their rate of serial order learning, based on participants’ performance in the case of repeated sequences of syllables. However, one account for this observation is that for many of the participants, spans for the infrequent syllables in fact included only few items, and hence - perhaps did not allow an adequate opportunity of sequence learning. In Experiment 2 of the current study, we use a protocol specifically designed for assessing learning of repeated sequences.

An elegant paradigm, specifically designed to assess sensitivity to repetition of the serial order is the Hebb repetition paradigm (Hebb, 1961). This is a sequence repetition task in which all the sequences are composed of the same items, but only one sequence is repeated in the same order (every third trial). Participants’ recall of the repeated sequence improves significantly compared with their recall of the non-repeated sequences, indicating specific benefits from sequence repetition (*The Hebb repetition effect*, Hebb, 1961). Since all sequences are composed of the same items, this paradigm allows to specifically track the rate of learning a repeated sequence (though note Siegelman, Bogaerts, & Frost, 2016). Importantly, the number of sequences administered (30) is larger than in Digit Span and does not depend on participant’s performance, and the scoring is per item rather than per sequence, hence providing more information.

The Hebb repetition learning paradigm was administered to adult IDDs in previous studies with mixed results. Initial studies reported reduced benefits from serial repetition (Bogaerts, Szmalec, Hachmann, Page, & Duyck, 2015; Szmalec, Loncke, Page, & Duyck, 2011) of sequences of Consonant-Vowel syllables presented auditorily, visually, and for spatial configurations of dots, in contrast to findings reported in Kimel et al. (Kimel et al., 2019). However, a follow-up replication study (Staels & Van den Broeck, 2015) did not reproduce this effect. Administering the same tasks to both children and adults, IDDs and adequate readers, they found no deficit in IDDs’ Hebb repetition effect, neither between the two groups of children, nor between the two groups of adults. Another study, in which adult IDDs and controls participated in a Hebb learning paradigm with short non-words, found a tendency of IDDs to have reduced serial learning, though the effect was not significant (Henderson & Warmington, 2017).

We now predicted that IDDs’ difficulties will not be with sequence repetition but with long-term frequency, and therefore tested Hebrew speaking participants (almost overlapping groups to those tested in Experiment 1, see Methods) on two versions of Hebb repetition task, using high and low frequency syllables.

We focused on syllables since (in Hebrew) they typically do not have a semantic component (i.e. most words are disyllabic), and exposure to them does not (strongly) depend on reading experience. Syllables’ presence as separate mental entities is evident quite early in development, whereas phonological awareness at the phonemic level requires explicit instruction and is reciprocally related to literacy (e.g. Perfetti, Beck, Bell, & Hughes, 1987). Indeed, illiterate adults and nursery-school children are similarly successful in manipulating syllables, but only literate individuals successfully operate with phonemes (Liberman, Shankweiler, Fischer, & Carter, 1974; Morais, Cary, Alegria, & Bertelson, 1979). Syllables’ unique psychological reality is kept in reading, where syllables probably form the basic automatically extracted unit (Ashby & Rayner, 2004; Carreiras, Alvarez, & De Vega, 1993; Prinzmetal, Treiman, & Rho, 1986).

## Method

### Participants

Twenty-five IDDs and 33 controls (vast majority of the pool described above; Tables 1-2) participated in the Hebb repetition learning experiment. Three control participants were removed from the analysis: one due to very poor performance in the Consonant-Vowel (open) syllables (CV) part (more than 4 SDs below the group mean) and two in order to match age with the IDD group (after matching: controls mean age(SD): 25.3(2.4), IDDs: 24(2.8)). Thus, the results for Experiment 2 are reported for 25 IDDs and 30 controls.

### Choice of frequent and infrequent syllables

We chose to use Consonant-Vowel (CV) syllables as our frequent items and Vowel-Consonant (VC) syllables, as our infrequent items. CV syllables are part of the syllabic vocabulary in almost all known languages (Sommer, 1970), and constitute the majority of syllables in Hebrew (Ben-Dror, Frost, & Bentin, 1995). Contrary to CV syllables, VC syllables are rare in languages in general (Clements & Keyser, 1983), and in Hebrew in particular (Ben-Dror et al., 1995). Converging evidence indicates that in Hebrew, CV syllables are an order of magnitude more frequent than VC syllables (Ben-Dror et al., 1995; discussed in Share & Blum, 2005). We verified this assumption by calculating the distribution of syllables in the largest publicly available punctuated corpus of spoken Hebrew (Kimel et al., 2019).

### Procedure

We administered the Hebb repetition learning task, in which sequences of nine syllables were presented to the participants in each recall trial; the same syllables were used for all sequences (Szmalec et al., 2011). The protocol was administered twice, once with CV and once with VC syllables, in a counterbalanced order across participants. The syllables were digitally recorded by a female speaker, and were presented to the participants through headphones. In each part we administered 30 sequences: 20 non-repeated sequences (“filler sequences”), and 10 repetitions of one sequence (the “Hebb sequence”). The repeated sequence was presented on every third trial, and its position (first/second/third) with respect to the beginning of the paradigm was randomized across participants. The played sequences were prepared in advance by randomizing sequences of syllables, and thus it is unlikely that a frequent bi or tri-syllabic sequence was presented consistently across participants (35 different sets of 20 non-repeated and 1 repeated sequences were used).

The paradigm was as follows: in each trial a pre-recorded sequence was presented and the participant was asked to orally reproduce the sequence, paying special attention to the order. If a participant knew that an item was missing from the recall, the participant was asked to verbally indicate it (e.g. *“ve tzu <u>pass</u> pi…*”). The score of each sequence was the proportion of syllables that were correctly reproduced with respect to their absolute position (e.g. if the participant recalled 2 syllables in their correct position, the score for that sequence will be 2/9 ∼= .22).

In order to study the dynamics of learning the repeated sequence, two regression lines were fitted to each participant’s scores across trials, one for the repeated sequence (10 repetitions) and one for the non-repeated sequences (20 sequences – the average of each 2 consecutive sequences was treated as one data point for the regression). The difference in the slopes of these two regression lines was used as a measure of serial order learning (after Szmalec et al., 2011). Given the reduced reliability of slope effects (Siegelman et al., 2016), we also assessed *Repetition, Syllable Type* and *Group* effects (and their interactions) using mean scores of each group under each condition: CV and VC for repeated and non-repeated sequences.

### Statistical analysis

The main analysis of means and slopes was done using a mixed-design ANOVA. In questions for which it was important to show that there was no group difference, we used the Bayesian approach, and reported a Bayes factor. Bayes factor compares the probability of the observed data (i.e., results of the experiment) given the null hypothesis (“the groups do not differ”) versus the probability of the observed data given the alternative hypothesis (“the groups differ”), thus allowing to compare the two hypotheses. Thus, unlike traditional frequentist statistics, it can provide support for the null hypothesis, and not just accept/reject the alternative. We used a calculator kindly provided by the *Perception and Cognition Lab* (Rouder, Speckman, Sun, Morey, & Iverson, 2009). It is generally accepted that Bayes factor values which are less than 3 are considered to support the null hypothesis (3-10 is some support, 10-30 strong, and greater than 30 very strong).

## Results

Overall, the results of the Hebb repetition paradigm were consistent with our hypothesis. Namely, IDDs had adequate benefits from series repetition, indicating adequate serial order learning, both when assessed with the slope and with mean performance. However, their overall scores were lower than controls’ when high-frequency syllables (CV) were used, but not when low-frequency syllables (VC) were used.

### Analysis of mean scores

We used a mixed-design ANOVA with *Syllable Type* (frequent vs. infrequent) and *Repetition* (repeated vs. non-repeated sequences) as within-subject factors, and *Group* (controls vs. IDDs) as a between-subject factor. We got the expected main effects: scores of frequent syllables were higher than scores of infrequent syllables (main effect of *Syllable Type*: *F*_(1,53)_ = 86.47, *p <* .001, *η2 =*.620; Figure 2; Table 4); scores of the repeated sequence were higher than scores of the non-repeated sequences (main effect of *Repetition*: *F*_(1,53)_ = 63.99, *p <* 10_-9_, *η2 =*.547; Figure 3; Table 4); controls had higher scores than IDDs (main effect of *Group*: *F*_(1,53)_ = 5.43, *p <* .024, *η2 =*.093 ; Figure 3A; Table 4).

**Table 4.** Experiment 2: scores (mean (SD)) of Hebb repetition learning with frequent and rare syllables. The score for each sequence was between 0 and 1: 0 if none of the 9 syllables composing the sequence was recalled correctly with respect to both its identity and serial position, and ∼.11 for each correctly recalled syllable. The total score for each subject for each condition is the average of the sequences’ scores.

| Type of syllable | Type of sequence | Controls | IDDs |
| --- | --- | --- | --- |
| Frequent | Non-repeated | .32 (.07) | .24 (.10) |
|  | Repeated | .50 (.20) | .39 (.18) |
|  | **Average repeated and non-repeated** | **.41** | **.31** |
| Infrequent | Non-repeated | .19 (.06) | .17 (.07) |
|  | Repeated | .31 (.18) | .26 (.14) |
|  | **Average repeated and non-repeated** | **.25** | **.21** |
| **Overall average** |  | **.33** | **.26** |

**Figure 2.**
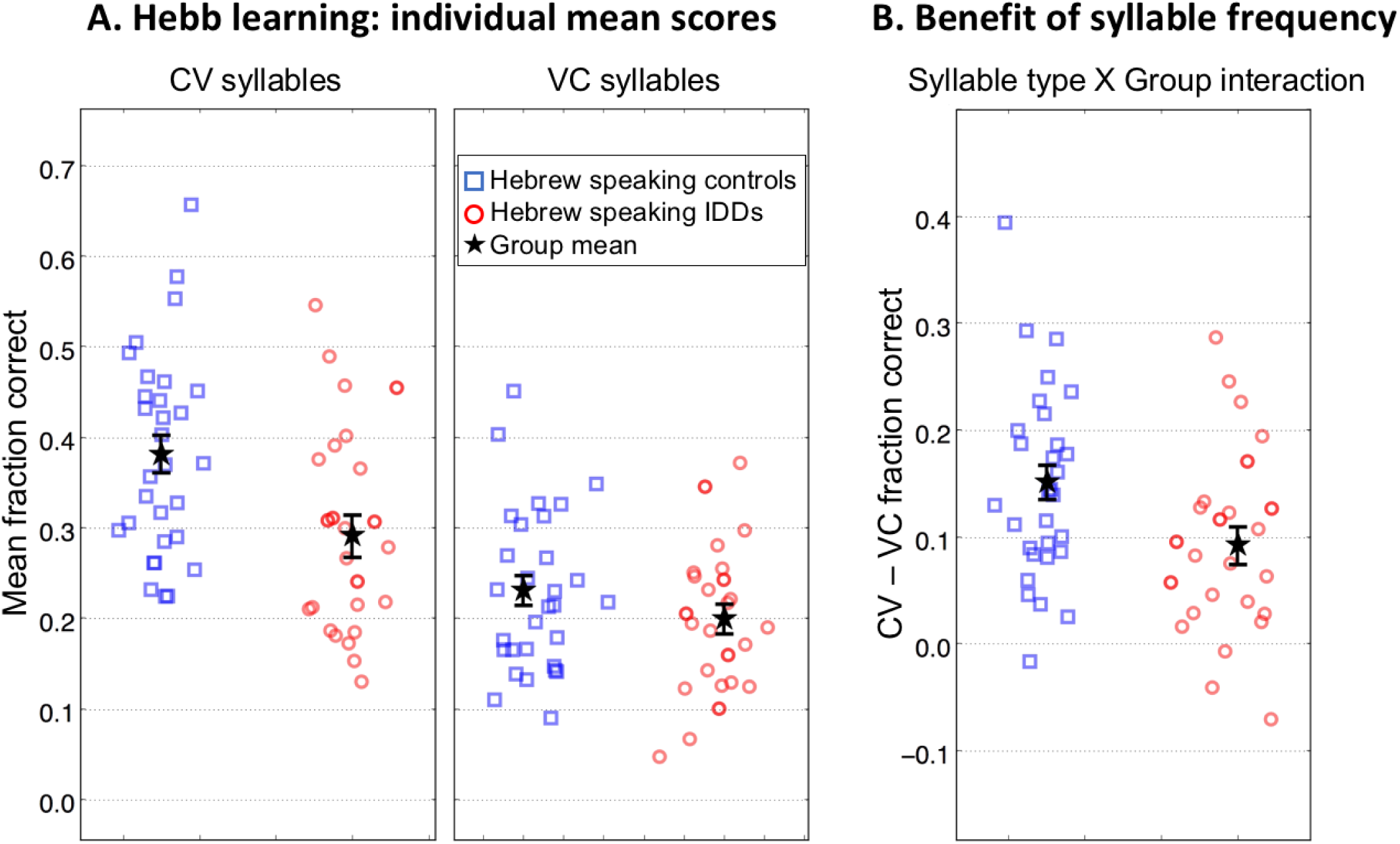
Hebb repetition learning paradigm: sensitivity to order using frequent (CV) and infrequent (VC) syllables: Hebrew speakers, controls versus IDDs. A. Mean scores across all sequences in the Hebb learning protocol of controls (blue squares) and IDDs (red circles). Left: frequent (CV) syllables. Right: infrequent (VC) syllables B. Benefits in spans (score differences) of CV compared with VC (long-term frequency effect). Controls gained more than IDDs from the use of the frequent CV syllables. Each symbol denotes a single subject’s mean score across all 30 sequences of the Hebb learning protocol. Error bars denote 1 SEM.

**Figure 3.**
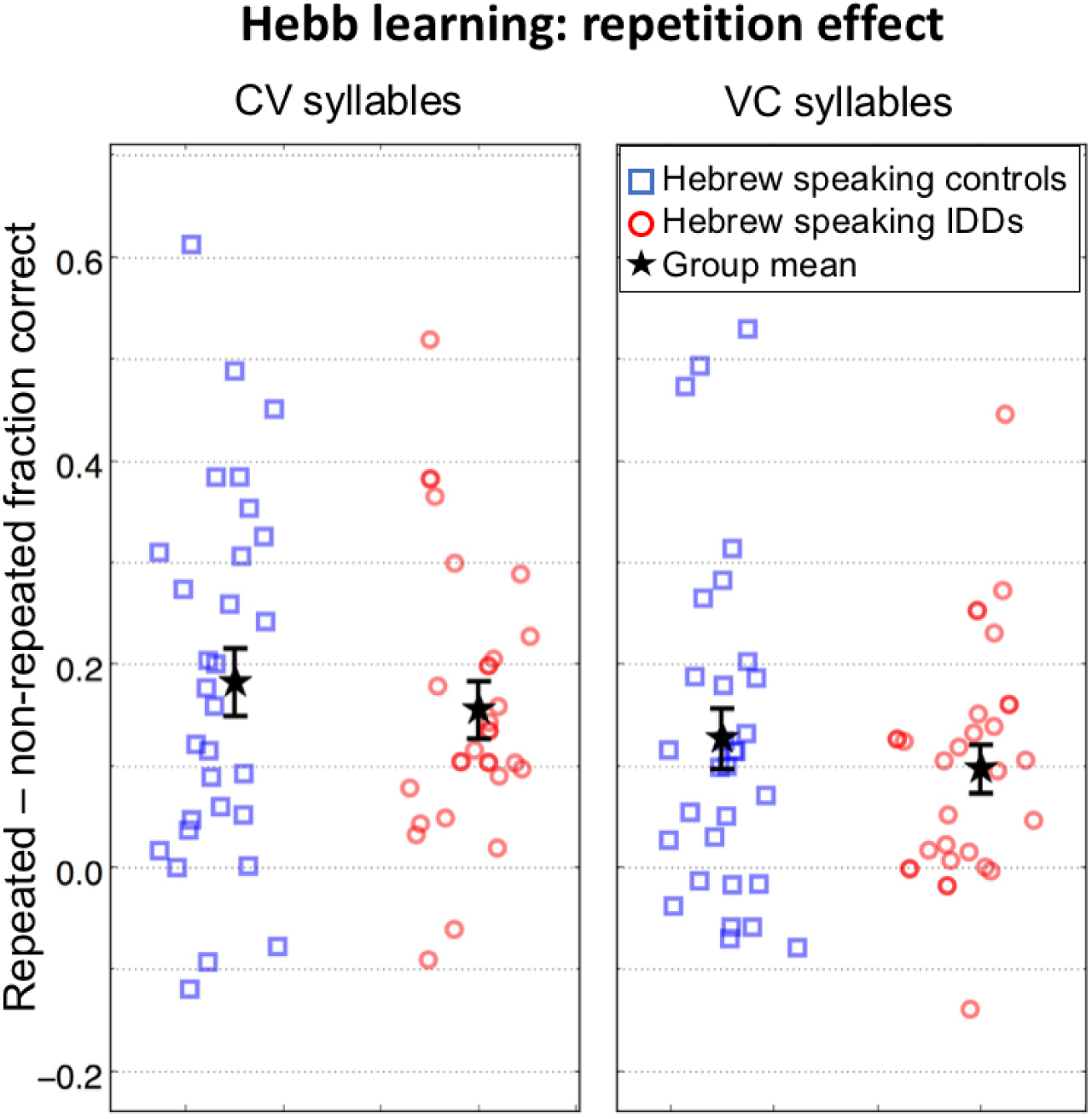
Difference between scores for repeated (Hebb) vs. non-repeated (filler) sequences of CV (left) and VC (right) syllables. The benefit is larger for CV syllables and is similar for controls (blue squares) and IDDs (red circles). Symbols denote individual difference scores. Error bars denote 1 SEM. The values are slightly jittered for display purposes.

Both groups significantly benefited from syllable frequency (controls: *d =*.160, *SE =*.02, *p <* 10_-10_; IDDs: *d =*.101, *SE =*.02, *p <* 10_-5_). However, importantly, controls benefited more than IDDs (*Syllable Type* X *Group*: *F*_(1,53)_ = 4.37, *p <* .041, *η2 =*.076; Table 4; Figure 2B). As expected, there was no significant group difference in the repetition effect (*Repetition* X *Group*: *F*_(1,53)_ = .64, *p =* .427, *η2 =*.012). The lack of difference between the groups (“null” hypothesis) is supported by the value of the Bayes factor: Scaled JZS Bayes Factor = 2.808, Scaled-Information Bayes Factor = 2.106, calculated on the benefit from repetition (Rouder et al., 2009).

Though repetition effect was significant for both the frequent CV syllables (*d =*.168, *SE =*.02, *p <* 10_-9_; Figure 3 - left) and the infrequent VC syllables (*d =*.111, *SE =*.02, *p <* 10_-6_ Figure 3 - right), the benefit from repetition was larger for frequent (CV) than for infrequent (VC) syllables (*Repetition* X *Syllable Type*: *F*_(1,53)_ *=* 6.01, *p <* .018, *η2 =* .102; Figure 3). There was no significant group difference in this repetition benefit for frequent compared with infrequent syllables (*Repetition X Syllable Type X Group*: *F*_(1,53)_ = .003, *p =* .958, *η*_*2*_ *<* 10_-4_; Figure 3).

### Analysis of learning rates

The Hebb learning paradigm (Hebb, 1961) also allows an assessment of learning rates by comparing slopes of the regression lines based on performance with repeated and with non-repeated sequences as a function of trial number for each individual (Szmalec et al., 2011). This analysis ignores the baseline performance, and incorporates only the learning rate. We calculated a regression line for each condition for each individual; the standard errors of the estimated gradients were small, and did not exceed .005 for any regression line, suggesting that the quality of the fit was satisfactory.

Learning rates for frequent syllables were higher than those for infrequent syllables (main effect of *Syllable Type*: *F*_(1,53)_ = 8.62, *p <* .005, *η2 =*.140; Figure 4 – top vs. bottom; Table 5), with no significant group difference (*Group* X *Syllable Type*: *F*_(1,53)_ = .01, *p =* .942, *η2 =*10_-4_; Figure 4 – left vs. right; Table 5). Learning rates of repeated sequences were higher than of non-repeated ones, as predicted (main effect of *Repetition*: *F*_(1,53)_ = 36.21, *p <* 10_-6_, *η2 =*.406; Table 5). Importantly, there was no significant group difference in the effect of sequence repetition on the learning rate (*Group* X *Repetition*: *F*_(1,53)_ = .01, *p =* .913, *η2 <* .001). The lack of difference between the groups (“null” hypothesis) is supported by the value of the Bayes factor: Scaled JZS Bayes Factor = 3.646, Scaled-Information Bayes Factor = 2.781, calculated on the benefit from repetition (Rouder et al., 2009). The effect of repetition on the learning rate was more beneficial for the frequent syllables than for the infrequent ones (Figure 4; *Repetition* X *Syllable Type*: *F*_(1,53)_ = 4.27, *p <* .044, *η2 =*.075; frequent: *d =*.024, *SE =*.004, *p <* 10_-7_, infrequent: *d =*.014, *SE =*.004, *p <* .002), with no significant group difference. Namely, there was no significant group difference in the repetition benefit for frequent compared with infrequent syllables for learning rate (Figure 3; *Repetition X Group X Syllable Type*: *F*_(1,53)_ = .03, *p =* .865, *η2 <* .001). In general, there was no significant group difference in the overall learning rate throughout the experiment (Figure 4, main effect of *Group F*_(1,53)_ = 2.73, *p =* .104, *η2 =*.049; Table 5).

**Table 5.** Experiment 2: slopes (mean (SD)) of Hebb repetition learning with frequent and rare syllables. The slopes of the regression lines were calculated for each individual based in each condition based on performance as a function of trial number.

| **Type of syllable** | **Type of sequence** | **Controls** | **IDDs** |
| --- | --- | --- | --- |
| **Frequent** | Non-repeated | .005 (.02) | -.00 (.01) |
|  | Repeated | .030 (.03) | .023 (.02) |
|  | **Average repeated and non-repeated** | **.017** | **.011** |
| **Infrequent** | Non-repeated | .004 (.01) | -.003 (.01) |
|  | Repeated | .017 (.03) | .011 (.03) |
|  | **Average repeated and non-repeated** | **.010** | **.004** |
| **Overall average** |  | **.014** | **.008** |

**Figure 4.**
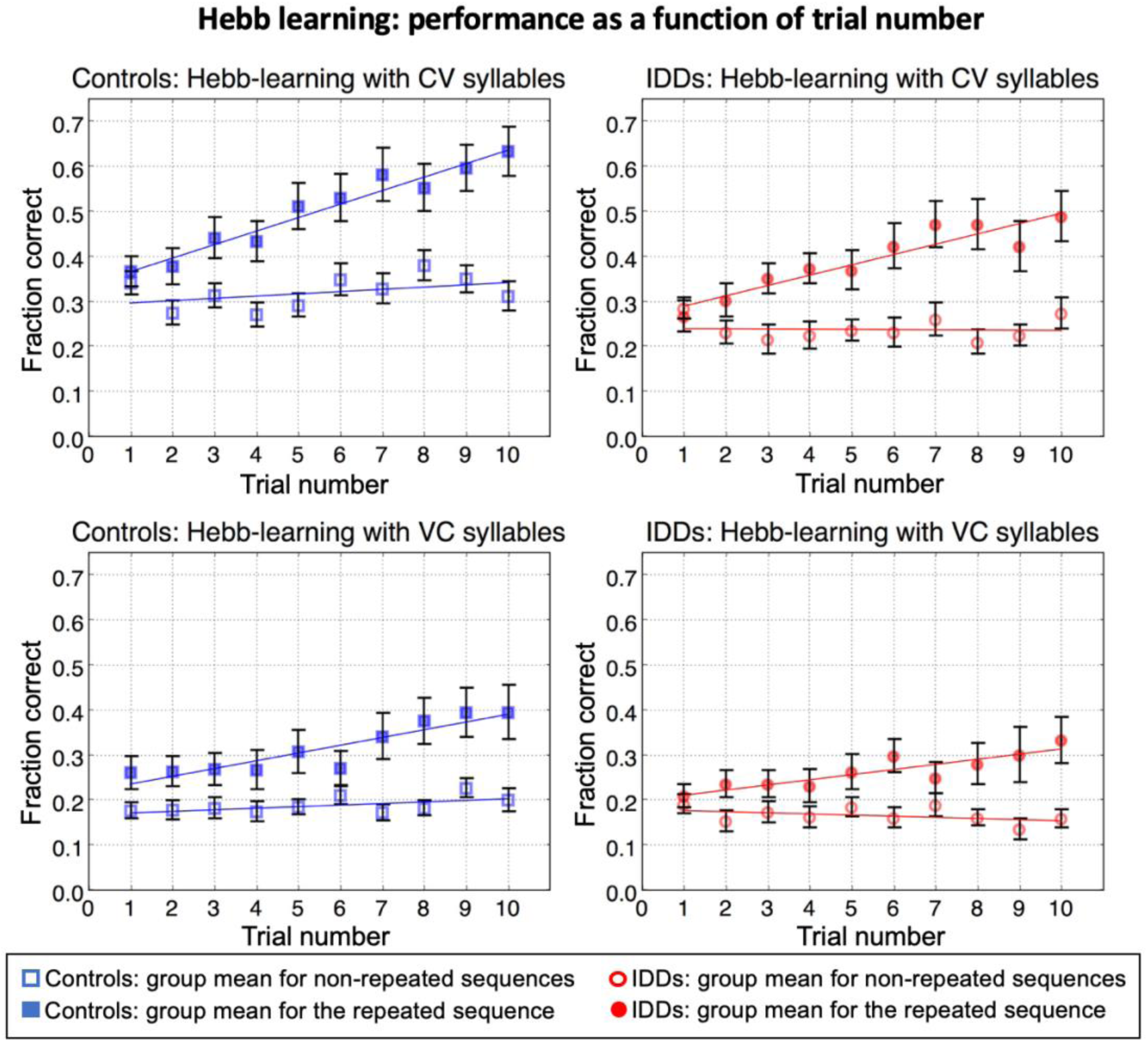
Fraction correct as a function of trial number for both repeated and non-repeated sequences of controls (left) and IDDs (right), for CV (top) and VC (bottom) syllables. Though controls’ CV scores for the non-repeated sequences are higher than IDDs’ (the regression line on the left has a higher intercept than the one on the right), the repetition-learning slopes do not differ between the groups. Error bars denote 1 SEM.

To summarize, we found a main effect of *Repetition*: slopes were steeper for the repeated vs. non-repeated sequences, and a main effect of *Syllable Type*, i.e. repetition effects were larger for CV than for VC sequences. There was also a *Repetition* X *Syllable Type* interaction. Namely, repetition slopes were steeper for CV than for VC syllables. However, the IDD and control groups had similar learning slopes, as shown in Figure 4, indicating similar rates of learning the serial order.

## Summary and Conclusions

The results of Experiment 2 were in line with the results of Experiment 1: both groups benefited from syllable frequency (e.g. Gathercole, 1995), but IDDs’ benefit was smaller than controls’. As for Hebb learning (the benefit of sequence repetition), IDDs’ benefits from sequence repetition did not significantly differ from controls’, both for frequent CV and infrequent VC syllables. Learning slopes were higher for CV compared with VC syllables, but still there was a significant learning also for repetition of VC syllable sequences. The slopes did not differ significantly between the groups. Importantly, all sequences (repeated and non-repeated) were constructed from the same syllables, and thus, the effects cannot be attributed to exposure difference within the experiment. In summary, Experiment 2 implies that IDDs’ long-term benefits from syllable frequency are smaller than controls’. Second, IDDs’ benefits from recent sequence repetition are not impaired; if there is any deficit in serial learning in dyslexia, it is small and not robust across studies and populations of IDDs (Majerus & Cowan, 2016).

## Concluding Discussion

### Summary of results

This study is composed of two experiments in which we assessed the effect of long-term item frequency and serial order learning on the capacity of STM in good readers and IDDs which were matched in age and cognitive skills. In Experiment 1, we found that controls benefit more from using native language items when tested with the common standard Digit Span test. In Experiment 2, using the Hebb repetition learning protocol, designed especially for assessing learning of serial order (Hebb, 1961), we found that controls benefited more than IDDs from syllable frequency, but the two groups did not significantly differ in their benefit from sequence repetition. Taken together these results suggest than the common observation of poor STM in dyslexia may largely reflect poorer item-specific long-term learning. The dissociation between poor sensitivity to items’ long-term frequency and adequate serial order learning suggests that different mechanisms underlie learning of single item long-term distributions and learning of repeated sequences. Only the former seems to be impaired in dyslexia.

### The profile of statistical sensitivities in dyslexia

We found that IDDs’ benefit from the long-term item frequency is reduced. These results are consistent with the anchoring deficit hypothesis (Ahissar et al., 2006) and with other theories of dyslexia that emphasize the a-typicality of the learning process in dyslexia. One such hypothesis is a core deficit in procedural learning (Lum, Ullman, & Conti-Ramsden, 2013; Ullman, 2004). It suggests a domain general learning problem which hinders skill automatization (Roderick I. Nicolson & Fawcett, 2007). Despite being domain general, the declarative-procedural distinction was also elegantly associated with language abilities (Ullman, 2004). The reduced benefit of IDDs from long-term frequency is in line with these accounts, though the exact nature of automatization in implicit syllable learning throughout life is yet to be specified.

Unlike theories of dyslexia in which the learning process is central and is used to account for failed skill acquisition, the phonological deficit account (Vellutino, Fletcher, Snowling, & Scanlon, 2004) and external (Sperling, Lu, Manis, & Seidenberg, 2005) or internal general noise accounts (Ziegler, Pech-Georgel, George, & Lorenzi, 2009) are less likely to explain the results of this study. More exposures allow more learning opportunities, and that should reduce the effect of noise for the frequent stimuli, or at least not increase it. Consequently, on the basis of these theories, we would expect group difference in spans using frequent items to be, at least, no greater than group difference in spans using infrequent items, whereas this study shows the opposite. Overall, our findings suggest that differences measured in phonological memory tasks are largely affected by long-term memory, consistent with findings indicating that performance in the span task is mediated by long-term knowledge (e.g. Gathercole, 1995). Particularly revealing is the lack of correlation among controls between Digit Span in Hebrew and English. Had there been individuals with generally good versus generally poor phonological short-term memory skills, regardless of their item-specific experience, we would expect a significant correlation within the control group too.

IDDs’ reduced benefit from long-term statistics for STM span tasks may stem from a number of reasons. The mental syllabary construct theory (Wheeldon & Levelt, 1994) suggests a different route of production for low vs. high frequency syllables. It suggests that the motor-programs for low-frequency syllables are constructed online from smaller components, whereas programs for high-frequency syllables are stored with a ready-to-use motor program. According to this account, IDDs’ deficit yields difficulties in the gradual formation of motor programs for repeating sequences, or in the efficiency of their use.

However, a study by Perez and colleagues (Perez, Majerus, Mahot, & Poncelet, 2012) suggests that the deficit is unrelated to production. In this study, participants were presented auditorily with a series of names of animals, and then had to order cards with pictures of these animals. This task was considered to tap only recall of order, since no recall of the items themselves was needed as they were given all the cards that they need, and only them. Children with dyslexia had reduced performance in this task compared to age matched controls and also compared to reading-level matched children. Importantly, only highly frequent and familiar animal names were used in the task, and thus the group difference in performance might reflect the reduced utilization of long-term item frequency in dyslexia. This deficit is not related to word production, but rather to general cognitive load since even just manipulating the order of highly familiar items is easier. According to the results of experiments 1 and 2 of the current study, if less familiar items were used in the order reconstruction task, overall performance would have been lower, but group difference would have been smaller.

The pattern of results in Experiment 2 did not replicate some earlier observations (Bogaerts et al., 2015; Szmalec et al., 2011), which found smaller slopes in IDDs’ Hebb learning and memory for sequences in other STM tasks (Oganian & Ahissar, 2012) but is consistent with subsequent studies (Henderson & Warmington, 2017; Kimel et al., 2019; Staels & Van den Broeck, 2014, 2015; Wang, Xuan, & Jarrold, 2016), which found no significant deficiency in IDDs’ Hebb learning or memory for repeated serial information. Thus, 6 studies did not find a significant deficit in repeated order learning, while 3 report a significant deficit.

Taken together, we conclude that if there is an effect, its size is small. We cannot, however, refute the account that there is a small effect of impaired learning of serial order. A significant effect was reported by two different research groups (Duyck et al., 2014; Oganian & Ahissar, 2012), and an insignificant trend in this direction can be seen in other studies (e.g., the current study, Henderson & Warmington, 2017; Staels & Van den Broeck, 2015).

### Challenging the concept of reduced STM in dyslexia

Though word and non-word spans are routinely assessed in dyslexia studies as a measure for STM capacity, most studies of dyslexia never considered the frequency of items which are used in these tasks. We are unaware of any previous study that compared auditory spans for frequent vs. infrequent syllables with dyslexic participants. The closest we are familiar with is one study that administered a non-word repetition task to children with and without dyslexia and manipulated phonotactic probability (Rispens, Baker, & Duinmeijera, 2015). Its results seem inconsistent with ours: children with dyslexia had greater relative difficulties with low phonotactic probabilities than with high phonotactic probabilities. However, this difference may stem from phonotactics not being the equivalent measure of syllabic frequency: the phonotactic probability (PP) in their experiment was based on bi-phones, which ignored syllable boundaries, and might have introduced a confound of syllable frequency in the “low-PP” and “high-PP” conditions. This is especially important since syllables are considered more natural units of processing than single phonemes or sequences of phonemes across word boundaries (e.g. Liberman et al., 1974). For example, in a study that used a masked priming paradigm, a string of letters was presented before a word or a non-word. When this string formed the first syllable of the target word, naming time was reduced compared with the case that this string contained an extra or a missing letter from the first syllable of the target (Ferrand, Segui, & Grainger, 1996). Age may also play a role in the inter-study difference. Specifically, the performance of a group of younger typically developing children showed a similar pattern of performance to the group of children with dyslexia, suggesting that children with dyslexia benefit less from exposure, making their performance similar to that of less experienced (younger) individuals (see also Kimel et al., 2019 for comparison of adult IDDs to non-native speakers).

Several studies administered spans of words, non-words and syllables, to reading-impaired children, and found reduced performance as compared to the control group. However, when words were used, lexical or semantic effects are likely to be relevant, and this might explain why a reduced word frequency effect was not found for IDDs (Davies et al., 2007; Van der Leij & Van Daal, 1999). When syllables were used, their frequency was not addressed (Brady, Poggie, & Rapala, 1989; Kamhi, Catts, Mauer, Apel, & Gentry, 1988; Snowling, 1981; Snowling et al., 1986). To summarize, we suggest that the numerous reports of reduced STM capacity in dyslexia largely reflect the choice of frequent items for these tasks, thus hampering specifically IDDs’ performance, due to their poor utilization of long-term knowledge, namely item frequency.

### Dissociating long-term distributional statistics from sequence learning

Memorizing a sequence of items introduces “sequence effects” which are not explained by item frequency alone. For example, when frequent words are presented as part of mixed sequences of frequent and infrequent words, they are remembered less well than when they are presented as part of a sequence of frequent words only (Hulme, Stuart, Brown, & Morin, 2003). The results of the Experiments 1 and 2 suggest that sequence learning in the course of repeated presentations does not strongly depend on single item frequency as well. Rather, it is supported by different mechanisms than sensitivity to distributional statistics, which underlies the benefit from very frequent language units. However, the benefits from sequence repetition were larger for CV than for VC sequences, observed in both groups. If sensitivity to serial order does not heavily depend on item familiarity, why should it be larger for CV compared for VC syllables? We attribute this difference to the inherent difference between CV and VC syllables, which does not directly relate to their frequency, but to the difference in the manner of their articulation, causing CV syllables to be more physically and physiologically “chunkable”. Based on their different type of ending (rime), CV syllables are categorized as “light” whereas VC syllables are categorized as “heavy” (Duanmu, 2010; Lunden, 2011). Heavy syllables attract stress (the Weight-Stress Principle, Duanmu, 2010; Velupillai, 2012) and a word can only have one primary stress. Therefore, a sequence of VCs tends to be articulated with a stress on each syllable, supporting segmentation rather than chunking, and perhaps impeding memory for the sequence as a whole, since stressed syllables are treated as word onsets (Cutler & Norris, 1988). By contrast, a sequence of CVs can be perceived and articulated as a long (chunked) multi-syllabic word.

Thus, we propose that controls benefit more than IDDs from the use of CV vs. VC syllables due to CVs’ higher frequency. However, controls do not benefit more than IDDs from sequence repetition since repetition effect does not depend on syllable frequency, but is mainly due to syllable “chunkability”, and the ability to chunk does not differ between the two groups. Both groups benefit more from sequence repetition of CV syllables vs. VC syllables since CV syllables are more chunkable than VC syllables due to mechanisms that are not exposure-dependent and do not differ between the groups. The separation between chunkability, which is not dependent on exposure and is not impaired in dyslexia, and item frequency for which sensitivity is reduced in dyslexia, was also found in a recent study (Kimel et al., 2019).

In summary, we measured the sensitivity of IDDs to two aspects of statistical learning, with different results for these two aspects. We found no deficit in IDDs’ learning of serial order. Benefit from sequence repetition did not differ between groups, in spite of recruiting a relatively large number of participants, and using a sensitive paradigm (Hebb repetition learning). Thus, if there is a serial order learning deficit in dyslexia, it is at most mild. By contrast, we found a deficit in the advantage of using high-frequency items compared with low-frequency items in each of the two experiments we conducted. In both experiments, controls and IDDs had larger spans for more frequently heard words and syllables. However, this benefit was significantly smaller in IDDs, indicating that their accumulative long-term benefit per exposure is reduced.

